# Iceberg or cut off – how adults who stutter articulate fluent-sounding utterances

**DOI:** 10.1101/2020.04.15.042432

**Authors:** Andreas Leha, Susanne Dickhut, Daniela Ponssen, Annika Primassin, Alexandra Korzeczek, Arun A. Joseph, Walter Paulus, Jens Frahm, Martin Sommer

**Affiliations:** Department of Medical Statistics, University Medical Center Göttingen, Germany; Department of Clinical Neurophysiology, University Medical Center Göttingen, Germany; Biomedizinische NMR, Max-Planck-Institut für biophysikalische Chemie, Göttingen, Germany

**Author notes:** **Correspondence:** Martin Sommer, M.D.

**Keywords:** Real-time magnetic resonance imaging, stuttering, articulation, trait and state markers

## Abstract

Whether fluent-sounding utterances of adults who stutter (AWS) are normally articulated is unclear. We asked 15 AWS and 17 matched adults who do not stutter (ANS) to utter the pseudoword “natscheitideut” 15 times in a 3 T MRI scanner while recording real-time MRI videos at 55 frames per per second in a mid-sagittal plane. All stuttered or otherwise dysfluent runs were discarded. We used sophisticated analyses to model the movement of the tip of the tongue, lips and velum.

We observed reproducible movement patterns of the inner and outer articulators which were similar in both groups. Speech duration was similar in both groups and decreased over repetitions, more so in ANS than in AWS. The variability of the movement patterns of tongue, lips and velum decreased over repetitions. The extent of variability decrease was similar in both groups. Across all participants, this repetition effect on movement variability for the lips and the tip of the tongue was less pronounced in severely as compared to mildly stuttering individuals.

We conclude that there is no major difference in the movement patterns of a fluent-sounding utterance in both groups. This encourages studies looking at state rather than trait markers of speech dysfluency.

## INTRODUCTION

The border between fluent and dysfluent speech is poorly charted and a no man’s land. Some rather fluent speaking people regard themselves as stuttering, other rather dysfluent speakers regard themselves as fluent. Stuttering is a speech fluency disorder that affects about 5% of children and about 1% of adults (Howell 2007). Symptoms comprise involuntary repetitions of syllables and sounds (e.g., “wi-wi-wizzard”), prolongations of sounds (“ffffffffffffree”) and tense pauses related to speech blocks (Bloodstein and Ratner 2008). Stuttering events may be accompanied by symptoms such as excessive effort also in muscles usually not involved in the speaking task, respiratory changes, or verbal and situational avoidance (Mulligan et al. 2001). Many affected individuals react with speech anxiety and feelings of shame (Iverach et al. 2009). Stuttering has a negative effect on social contacts, job opportunities and personal development (McAllister, Collier, and Shepstone 2012).

Cerebral neuroimaging methods have much improved over the last 20 years in visualizing brain structural and functional correlates of stuttering (Belyk, Kraft, and Brown 2015). By contrast, examining in detail what the articulators actually do during stuttered speech has been notoriously difficult in the past and has only slowly progressed over the last years. It requires methods with a high temporal resolution, such as electromyography. All of these, however, can only be used to a limited extent intraorally, and are limited by the recording electrodes themselves hindering the flow of speech (Platt and Basili 1973; McClean, Goldsmith, and Cerf 1984). Therefore, there are only few reliable laboratory settings, such as using optical markers or optomagnetic devices to record motion sequences, but these are mostly confined to the skin surface and lips, with only limited insight into inner speech organs (Smith et al. 2012).

This methodological and technical limitation has now been overcome by real-time magnetic resonance imaging (MRI), which is capable of visualizing the articulator’s motion sequence at 30 to 100 frames per second while simultaneously recording audible speech (Uecker *et al*., 2010). Using this novel MRI technology we performed a study investigating articulation movements in a group of adults who stutter (AWS), comparing the results to a group of adults who do not stutter (ANS).

We here probe the hypothesis that articulation in AWS is normal in fluent-sounding speech sections. The alternative hypothesis is that articulatory movements of AWS and ANS also differ in fluent-sounding speech sections. Indeed, it is unclear where fluent speech ends and where stuttering begins, i.e. whether the border between fluent speech and stuttering is a ‘gradual transient’ or an abrupt ‘all or none’ phenomenon. The gradual transient assumption would mean that stuttering is a continuous phenomenon audible only when reaching a certain threshold, a view which has been illustrated as an iceberg (Sheehan 1970), where the undoubtedly visible part is only a small fraction of the true phenomenon. The ‘all or none’ view implies that stuttering events are special, and fluent speech segments normal. This ongoing matter of debate is further blurred by “covert stuttering”, i.e. affected individuals hiding their speech impediment. Such individuals often develop highly skilled avoidance behavior, despite which many carry a high emotional burden, others are very dysfluent but not regard themselves as stuttering, at least not in public (Douglass, Schwab, and Alvarado 2018).

In childhood, developmental dysfluencies are distinct from early characteristics of persistent stuttering (Sandrieser and Schneider 2008). Hence, we here hypothesized an all or none phenomenon rather than a gradual transition between fluent and dysfluent speech. Hence, we expected the articulatory movements producing fluent sounding utterances of adults who stutter not to differ substantially from the movement patterns of utterances of the non-stuttering peers. We chose the fluent sounding runs of the pseudoword “natscheitideut” from material obtained in an M.D. thesis (Ponssen in preparation). We aimed at studying movement patterns, variability, and possible learning effect over repetitions.

In detail, we hypothesized:

### Hypothesis 1

Across individuals, reproducible movement patterns of the articulators can be recognized when the pseudoword “natscheitideut” is uttered in an audibly fluent manner. In adults who stutter, the lip, tongue and velum movements during fluent-sounding speech are similar to the fluent speech of nonstuttering subjects.

### Hypothesis 2

The variability of the movement patterns of tongue, lips and velum decreases in the individual subjects over the number of repetitions of the pseudoword “natscheitideut”. This learning effect is present in both groups.

Smith et al. (2010) measured the lip opening variability of stuttering and non-stuttering volunteers in fluently-spoken pseudowords of varying length and complexity. In the comparison of five early and five late runs, the word “mapshaytiedoib”, which is similar in complexity and length to the word “natscheitideut” used in this study, showed a reduction of the variability of lip movements in both the stuttering and non-stuttering subjects the variability of lip movements. This was significant in the stuttering subjects but not significant in the non-stuttering subjects (Smith et al. 2010). We intended to confirm and expand this not only assessing the variability of the lip movement, but also the movement of the tongue and the velum movement.

### Hypothesis 3

There is a group difference in the decrease in the variability of movement patterns. Real-time MRI confirms findings from the recording of optical markers on the lips regarding less stable lip movement cycles in stuttering subjects compared to non-stuttering subjects during non-word repetitions. These are present not only in the lips, but also in the tongue and the velum. Stuttering subjects initially show more variable articulations than non-stuttering subjects. The learning effect, which is measured by a reduction in variability, is greater in stuttering subjects than in non-stuttering subjects (Smith et al. 2010).

### Hypothesis 4

Speech duration of the fluently spoken pseudoword “natscheitideut” is similar in both groups. Even after repeating the pseudoword several times, the time does not change.

### Hypothesis 5

The stuttering severity of the stuttering subjects accentuates and modulates the observed group difference. In the early repetitions, severely stuttering subjects have a high variability of articulation movements. In the later repetitions, however, they show a greater learning effect than less affected individuals.

## METHODS

The protocol was approved by the University Medical Center Göttingen ethics committee, and we obtained written informed consent before any study-related procedure took place.

### Participants

We investigated 15 subjects with persistent developmental stuttering (PDS). Clinical characteristics are shown in table 1. They were recruited from the “Institut der Kasseler Stottertherapie” (Euler et al. 2009) and the Göttingen stuttering peer support group. Their data was compared with those from 17 matched ANS with no personal or family history of stuttering. None of the participants had had any unstable medical or neurological prior illnesses, and none of them was taking CNS-active drugs at the time of participation.

**Table 1:**
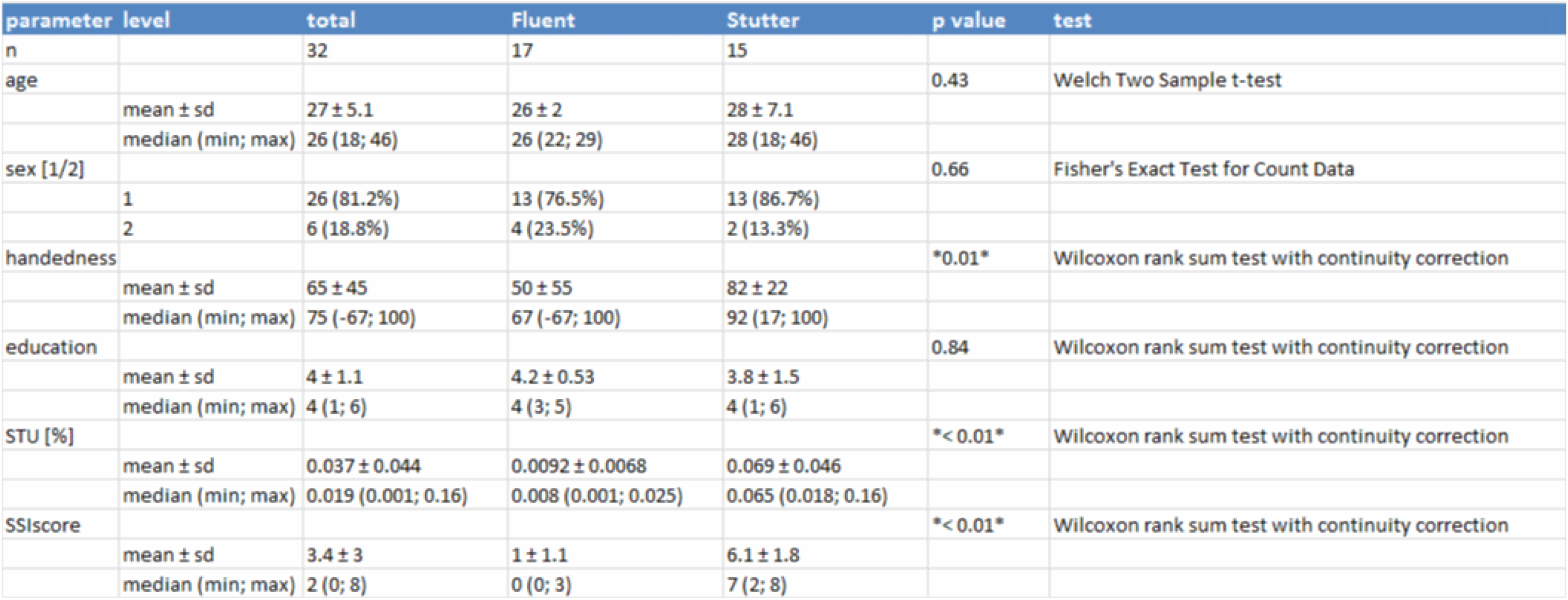
Epidemiologiical data of participants. Comparison of age, gender, body height, handedness and percent of stuttered syllables of all participants.

We obtained real-time MRI videos of 15 repetitions of a pseudoword. The first 10 fluently spoken recordings per individual were extracted. Repetitions with stuttering or mis-pronounciations were not eligible for analysis. Additionally, the first two recordings were regarded as training repetitions and not considered for analysis. The remaining repetitions were divided into two halves, an early phase and a late phase. As maximally 10 out of 15 repetitions were analysed, even with relatively strong stuttering subjects a sufficient number of completely fluently spoken runs could be evaluated.

#### SSI-IV

The severity of the stuttering of the individual subjects was assessed by the speech therapist Mrs. Bettina Helten with the help of the SSI-IV scores. The SSI-IV provides a reliable and valid norm-related assessment of the severity of stuttering. Speech samples are composed of a conversation and a text, which is read aloud by the subjects. The conversation should include motivating topics for the subject and the scope should be 150 to 300 syllables (Sandrieser and Schneider 2008).

### Choice of linguistic material

We chose pseudowords involving different articulation loci, hence expanding the existing literature which focusses on bilabials visible to external movement tracking (Smith et al. 2010). In this study, the pseudoword “natscheitideut” was chosen, which consists of the alveolar consonants [n], [d] and [t] and the postalveolar consonant [∫]. This focusses on tongue and velum movements well visible in real-time MRI videos in a mid-sagittal image orientation through the oral cavity. The articulation locations of the tip of the tongue on the anterior upper incisor and on the hard palate behind the upper incisors, which belong to the alveolar and postal convex consonants of this pseudoword, are thus represented. With such real-time MRI datasets, therefore, the relevant articulation positions of the pseudoword “natscheitideut” can be analyzed (Niebergall et al. 2013).

Fluent speech sections were defined by the absence of typical stuttering characteristics such as repetitions of sounds, sound expansions or blockages. In addition, we discarded longer pauses or hesitations between two words or within a word if they led to a pause of more than 250 ms (Van Zaalen-Op ’t Hof, Wijnen, and De Jonckere 2009). In addition, recordings were discarded if subjects made pronunciation errors (e.g. ‘natscheideut’ instead of ‘natscheitideut’).

### Real-time MRI

Technical details on real-time MRI are provided elsewhere (Uecker et al. 2010). In brief, participants were supine in a 3 T MRI system equipped with a 64-whannel head coil (Siemens Healthcare, Erlangen, Germany). Participants wore ear plugs and an MRI-compatible headphones connected to an MRI-compatible microphone (FOMRI III+, Optoacoustics Ltd, Israel). Real-time MRI was based on highly undersampled radial fast low-angle shot (FLASH) acquisitions. Recordings were in a mid-sagittal orientation covering the entire vocal tract, using an in-plane resolution of 1.4×1.4 mm^2^, a slice thickness of 8 mm, a field-of-view of 192×192 mm^2^ and a base resolution of 128 data samples per radial spoke. Acquisitions employed the following parameters: repetition time (TR) =2.02 ms, echo time (TE) =1.28 ms, flip angle =5°, 9 spokes, measuring time per image =18.18 ms corresponding to 55 frames per second.

Four pseudowords („gakscheitideuk“, „maptibibi“, „natscheitideut“ and „mapscheitideup“) were visually presented on an MR-compatible screen, 15 times each in pseudorandomized order, embedding each word in a German carrier phrase (“Sa**g** XXX **b**itte”).

### Film Clip Cutting

We extracted relevant sections from the individual real-time MRI videos using the programs Audacity (Audacity software copyright 1999-2019 Audacity Team) and VirtualDub (version 1.10.4, Avery Lee). As starting point and end point, the images corresponding to the two plosives [g] and [b] of the images “sa**g** natscheitideut **b**itte” were used.

With the open source program Audacity, it was possible to display the real-time MRI recordings as a spectrogram. The spectrogram shows three dimensions. The time is displayed on the x-axis, the frequency on the y-axis, and the color intensity indicates the energy level. In the spectrogram, the plosives and thus the start and end points ([g] and [b], see above) could be detected. A plosive is subdivided into the phase of closure formation in which, for example, the / g / of the back of the tongue migrates to the hard palate and closes the interspace, or in the case of / b / the closure of the upper and lower lip. In the closing phase, the expiratory air flow is stowed behind the back of the tongue or behind the lips. Over a short period of time, this condition is maintained. In the spectrogram you can see in these voiced vowels a so-called “voice bar” at below 500 Hz. The third phase is the sudden solution under explosive noise. The shutter sound takes only 10-20 ms for voiced plosives. The plosive is easy to recognize and to determine based on the closing phase, which can be seen on the “voice bar” and https://www.phonetik.uni-muenchen.de/studium/skripten/SGL/SGLKap2.html and (Pompino-Marschall 2009) (supplementary figure 1).

Hence, it is possible to exactly determine the starting point. As a control, the open source program VirtualDub was chosen. With this program it is possible to visually and audibly view the real-time MRI image series. The times determined with Audacity could be compared with the images of the real-time MRI. With VirtualDub it was possible to look at each picture individually. At the starting point, therefore, the back of the tongue had to be on the soft palate, and at the end point, the lip closure for the [b] had just been completed. Thus, possible errors of the determination with the spectrogram could be improved upon (supplementary figure 2).

### Processing of film clips

We further processed the recordings in the MATLAB R2017b program with Image Processing and Signal Processing Toolbox (The Mathworks, Inc., Natick, Massachusetts, USA), which was specially developed for the dynamic data analysis of real-time MRI videos (Iltis et al. 2015). The toolbox places a grid over an image of the real-time MRI video. All pixels on which the grid is located can now be viewed as a function of time. The grid consists of a baseline, which is drawn between two fixed points in the image. Here we used the upper front edge of the cervical vertebrae 4. As a second fixed point, the transition from the anterior incisor to the hard palate was chosen, and the baseline was drawn between these two fixed points. The midpoint of the baseline was the starting point for seven more lines drawn automatically at angles of 0, 30, 60, 90, 120, 150, and 180 degrees between the two halves of the baseline (supplementary figure 3). The length of each other line was half the baseline length. Now the pixel values of the individual lines of the grid were analyzed over the time of the real-time MRI scan, and a line profile was created showing the pixel resolution of a single grid line on the Y axis and the time change on the X axis. This way, each line profile shows the pixels of each gridline over time. These line profiles can be used for further evaluation (Iltis et al. 2015).

Niebergall et al. (2013) showed that in real-time MRI the articulation positions of the articulator tongue, lips, and velum can be well visualized when speaking consonants. The pseudoword “sag natscheitideut bitte”, which is to be evaluated, consists of the alveolar consonants [n], [d] and [t] and the postalveolar consonant [∫]. In addition, the velar consonant [g] and the bilabial consonant [b] of the two auxiliary words “sag” (“say”) and “bitte” (“please”) are part of the evaluation. In order to find out how to modify the grid in order to visualize relevant articulation movements, three images of stuttering and three non-stuttering subjects were extracted at the time of consonant pronunciation. Based on these images and in comparison with the results of Niebergall the grid has been changed. Starting from the baseline, three lines were drawn to obtain information about the tongue tip movement in the alveolar and postal convex consonants. Starting from the midpoint of the baseline to the hard palate, these three lines were spaced 0 degrees, 7.5 degrees, and 15 degrees. Information about the velar consonant [g] with the location of the articulation of the back of the tongue, which hits the hard palate, was expected to be found by a line spaced 60 degrees apart. Another 90-degree line reflected a point in the velum movement. The next line was a line between the center of the upper lip and the center of the lower lip of the subjects. In this way, the lip movement in the bilabial consonant [b] can be detected. The last line is the back of the baseline from the midpoint to the spine. It records no relevant articulation, but controls for head movements (see figure 1).

**Figure 1:**
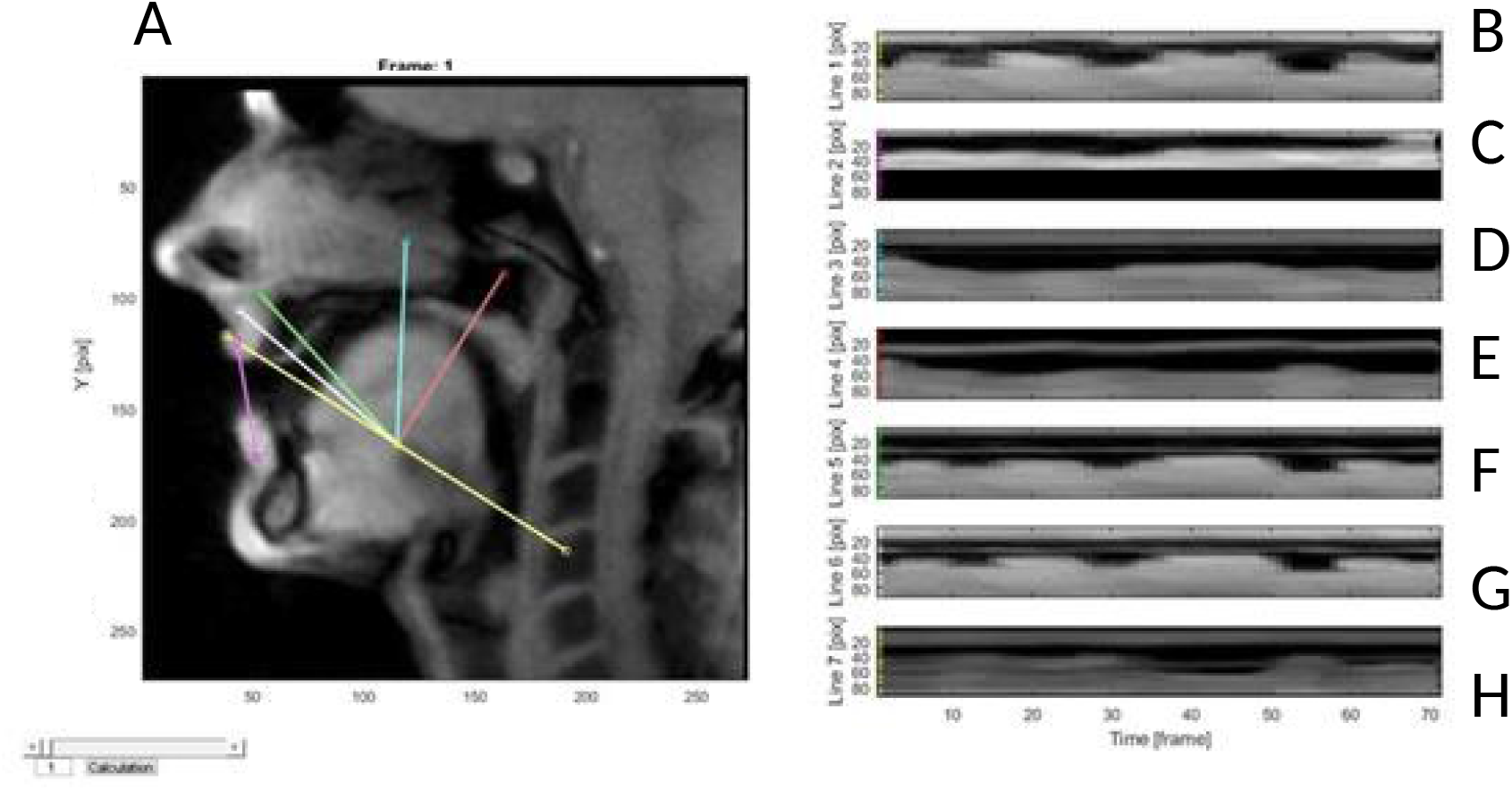
Movement analysis of articulators over time while speaking the pseudoword “natscheitideut”. Line profiles were adjusted to the first image, and the change of pixel contrast over time calculated. From this, the width of the main stretch of low-contrast (“black”) pixels over time was extracted and depicted here. The grid consists of a baseline, which is drawn between two fixed points in the image. Here we used the upper front edge of the cervical vertebrae 4 and the transition from the anterior incisor to the hard palate. The midpoint of the baseline was the starting point for seven more lines drawn automatically at angles of 30, 60, 90, 120, and 150 degrees between the two halves of the baseline. The length of each other line was half the baseline length. A, overview of first image with inserted grid, B-H, 7 derived time courses, covering the tip of the tongue (B, yellow line), the tongue and the anterior hard palate (C, purple line), the tongue and the posterior hard palate (D, blue line), posterior tongue and soft palate (E, orange line), posterior tongue and upper pharynx green line, and posterior tongue and middle pharynx, (G, white line), as well as baseline (H, second yellow line).

During the recording some subjects moved their head between the different pseudowords. For this reason, the grid was readjusted for each subject at every new repetition. The starting point for the grid adaptation was the first frame of the section to be analyzed with the articulation position of the consonant “g”.

### Processing of line profiles

Information had to be filtered from the line profiles to be able to compare the movements with each other. In MRI, the different tissues and airspaces are mapped as pixels in different shades of gray. The measurement of gap widths or distances from reference points in the line profiles was accomplished semi-automatically using in-house developed software. Matter was differentiated from air space based on the gray value of the pixels. Constrained three-component Gaussian mixture models were fit to the distribution of the gray values to classify matter into air space (dark), matter (light) and uncertain (gray) parts. The black component of the mixture distribution was used to set a threshold for image binarization. Detected gaps were registered over time by overlap to previous gap pixels. The gray-value threshold and the choice of gap to retain in case of multiple detected gaps were allowed to be manually overridden in an interactive application. Thus, the gaps can be accurately detected because the pixel values of the tongue, lip and hard palate clearly differ from the pixel value of the air space between the anatomical structures (supplementary figure 4).

Ann Smith compared the articulation movements of the lips using the lip-opening signal, which is calculated by subtracting the lower lip signal from the upper lip signal. Thus, the gap between the upper and lower lip was compared (Smith et al. 2010). In our analysis, for the most part, gaps are measured, which can then be analyzed and compared in a further step. In the line profiles to the alveolar and postalveolar articulation locations with the lines spaced 7.5 degrees and 15 degrees, the distance between the tip of the tongue and the hard palate is extracted. It is possible to take the gap as a measure of tongue movement because during articulation the palate does not move, but remains in the same position. Furthermore, the line profiles of the lines for bilateral consonants and for velar consonants were processed according to the same scheme, the space between the upper lip and lower lip or the space between the back of the tongue and the soft palate was extracted. With the line profile of the velum, it was not possible to take the space between the lower edge of the velum and the back of the tongue, because the back of the tongue also moves during articulation. The extracted space would not only represent the velum movement, but a combination of the movements of the velum and the back of the tongue. For this reason, the transition of the lower limit (liminal line) of the velum to the gap was chosen, and the spatial change of this transition was extracted over time. Also for the first line profile for the alveolar and postal convex consonants with the line which was the baseline, no gap could be analyzed. This is because the gray value of the pixels of the incisor is similar to that of the air. For many images, it is not possible to tell the difference between the back of the tooth and the air, and no precise gap can be defined. Therefore, the spatial movement of the transition of the tongue tip to the air space of the oral cavity was extracted, similar to the line profile for the velum (supplementary figure 5).

### Statistical analysis

Pretest results were compared between groups using unpaired, two-tailed t-test, Mann-Whitney-U test, or Fisher’s exact test, as appropriate.

To address hypothesis 1, the gap widths were scaled to the same duration and penalized flexible functional regression was applied to assess group effects in the time curves while accounting repeated measurement of each subject by adding subject as random time-varying effect. Modeled curves per subject as well as per group have been visualized.

To address hypotheses 2 and 3, smoothed curves were generated by approximation with (k = 15) cubic regression spline basis functions. The smoothed curves were centered around zero scaled to unit variance. For each subject the repetitions were grouped into two phases: an early phase and a late phase. The time was binned into 42 equally sized time windows and in each phase for each subject the standard deviation was calculated in each time window. The resulting standard deviations were summed up in each phase for each subject to arrive at a variation index.

In each place of articulation (lips, tip of tongue, back of tongue, velum) a linear mixed effect model was fit to the variation index with phase (early and late), group (ANS, and AWS) and their interaction as fixed effect and including the subject as random effect. Additionally, one model was fit across all places of articulation including the place of articulation and all interactions with it as additional fixed effect.

To address hypothesis 5, the percent of syllables stuttered from the reading and the spontaneous speech task were averaged into a speech fluency covariate. This speech fluency covariate was added to the linear mixed effect models of the variation index.

To address hypothesis 4, a linear mixed effect model with negative exponential repetition effect was fit to the durations to pronounce the word. The exponential repetition effect was chosen to account for longer durations in the first repetitions of pronouncing an unfamiliar pseudoword. Besides the negative exponential repetition, the group (ANS vs AWS) and their interaction were used as independent variables and the repeated measurements were accounted for by a random intercept per subject.

All analyses on the extracted images have been done in R (version 3.6.1; R Core Team 2018) using the packages EBImage (version 4.25.0; Pau et al. 2010) for image manipulation, shiny (version 1.3.2; Chang et al. 2019) to support the semi-automated gap detection, tidyfun (version 0.0.82; Scheipl, Goldsmith, and Wrobel 2020) and refund (version 0.1.21; Goldsmith et al. 2019) for the handling and modelling of the functional data, and lme4 (version 1.1.21; Bates et al. 2015) for the mixed effect modeling where p values were calculated via Satterthwaite’s degrees of freedom as implemented in lmerTest (version 3.1.0; Kuznetsova, Brockhoff, and Christensen 2017).

## RESULTS

Epidemiological data and the results of the pre-test questionnaires are listed in table 1.

The results of the SSI-IV show that the group of stuttering subjects consists of five subjects with very slight stuttering, three subjects with mild stuttering, four subjects with moderate stuttering, three subjects with severe stuttering, but no subject with very severe stuttering. In addition, the group of fluent subjects was assessed according to SSI-IV. Of the 17 subjects, 14 were found to not to be stuttering, while three subjects were classified as very mild stutterers.

To address <u>hypothesis 1 (reproducible movement patterns)</u>, we modelled movement patterns using subject and group, and assessed the group effect on the movement patterns. The patterns differed significantly between subjects but only slightly so and showed a high similarity across subjects (figure 2 A and B and supplement). Only in dimensions 2 and 4 the two groups exhibited slightly different movement patterns, while the modelled movement patterns are not significantly different between the groups in the remaining dimensions (figure 2 C). As a result, movement patterns were distinguishable but still very similar across subjects, and only slight differences between groups were observed. Hence, hypothesis one is confirmed for the overall sample without major group differences.

**Figure 2:**
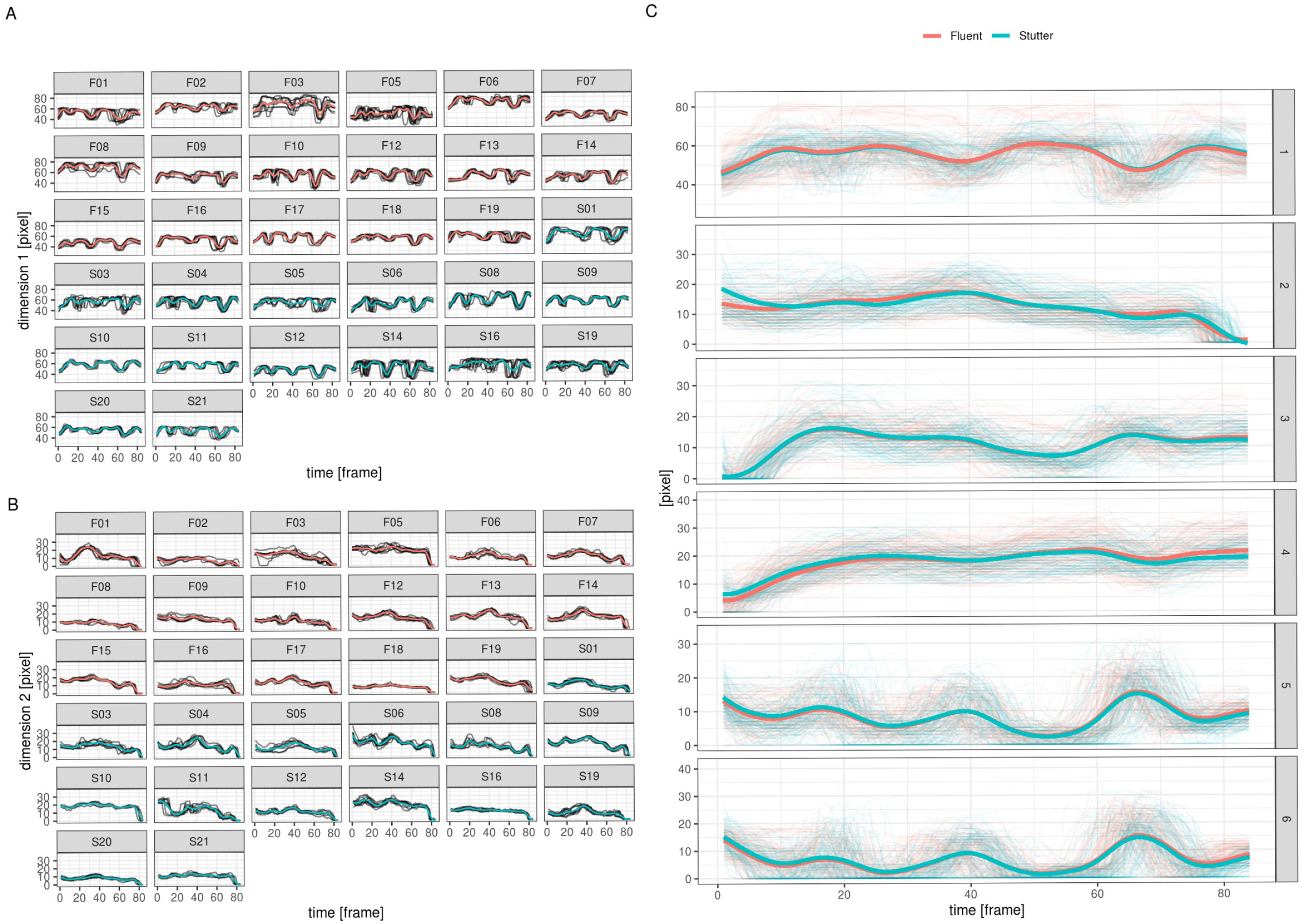
Modeled movement of articulators over time. The movement patterns were modeled using mixed-effect penalized functional regression models. (A and B) Individual model predcitions for dimensions 1 and 2; thin lines = measured individual profiles for each repetition, thick lines = model predictions for each subject. (C) Group-level model predictions for all dimensions; thin lines = measured individual profiles, thick lines = group level model predictions.

<u>Hypothesis 2</u> postulates that the variability of movement patterns decreases over time. Hy<u>pothesis 3</u> assumes a stronger decrease of variability over time in AWS than in ANS (Smith et al. 2010). To address these hypotheses we compared variability across articulators. We speculated that those with more degrees of freedom of movement in the sagittal plane would yield higher variability. Taking the tip of the tongue in the early half of the runs in fluent speakers (ANS) as reference, all other articulation loci (lips, back of the tongue, velum) differed significantly by showing less variability (figure 3 and supplementary table 1). We also observed a significant time (early vs. late) effect in that the variability was significantly lower in the second half. We did not observe a significant difference between groups (AWS vs ANS) and also the effects between articulation loci and time were not significantly different in the AWS group as the interactions were unrevealing (see supplementary table 1).

**Figure 3:**
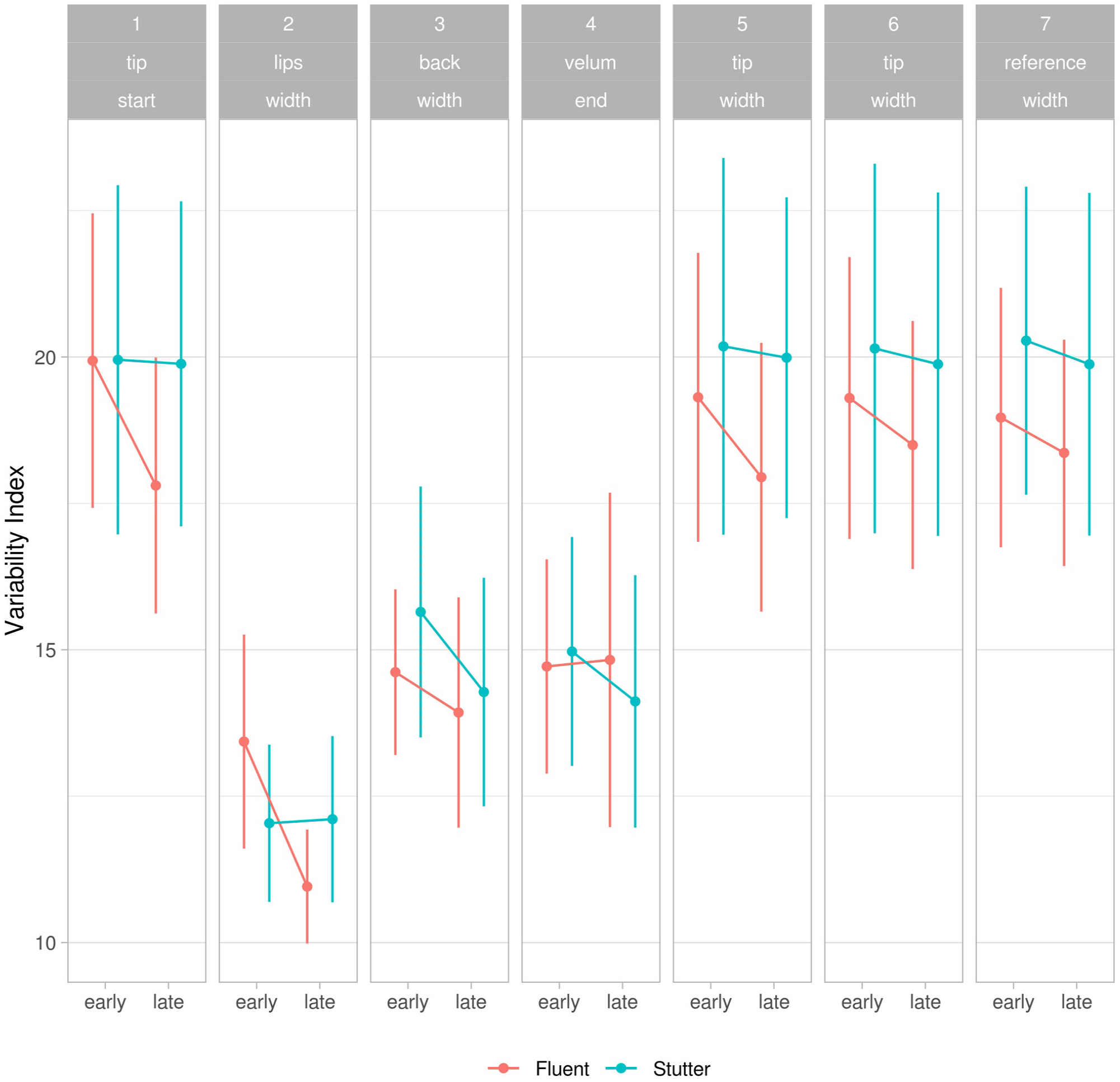
Variability index. Variability of articulatory movements for different articulators, in early and late trials, and in adults who stutter and a group of control participants. For details see text.

<u>Hypothesis 5</u> postulates that the stuttering severity of the stuttering subjects accentuates and modulates the observed group difference. To address this hypothesis, we included the stuttering severity as a covariate into the model (supplementary table 2). In the overall model stuttering severity (STU) did not exhibit a significant influence, and also the interactions were unrevealing.

As a second step, we compared movement variability of the early and the late phase of each articulator (figure 3). Regarding the analysis of each articulator separately, and looking at the variability index at the lips, late runs have a significantly lower variability than early runs (see supplementary table 3 A). Speech fluency tended to modulate this effect by lessening the time effect with increasing dysfluency.

The same type of analysis was used for the variability index at the tip of the tongue (see supplementary table 3 B): here, again the late runs show significantly less variability. This effect is absent or much smaller in the AWS group (see also figure 3), but this effect is not significant. The results for the variability index of the velum and the back of the tongue were unrevealing (see supplementary table 3 C and D).

Summarizing the findings on Hypothesis 3, we observe reduced variabiliy for the ANS group at the lips as well as the tip of the tongue (supplementary table 3 A and B). This reduction appears to be smaller in the AWS group (figure 3) with borderline significance at the lips (supplementary table 3 A). At the velum and the back of the tongue, there appears to be a higher reduction in the AWS group (figure 3) but the analytical results were unrevealing (supplementary table 3 C and D). We have an opposite pattern of dysfluencies on articulator variability.

<u>Hypothesis 4</u> assumes no difference overall regarding the timing of speaking. We addressed this by analyzing the duration of speaking of the entire utterance, defining the beginning as the pronounciation of the plosive [g] and the end as the pronounciation of the plosives [b] of the phrase “sa**g** natscheitideut **b**itte”, as illustrated in figure 1. This resulted in a significant interaction of group with time as the reduction in time was observed to be significantly larger in the AWS group compared to the ANS group (figure 4).

**Figure 4:**
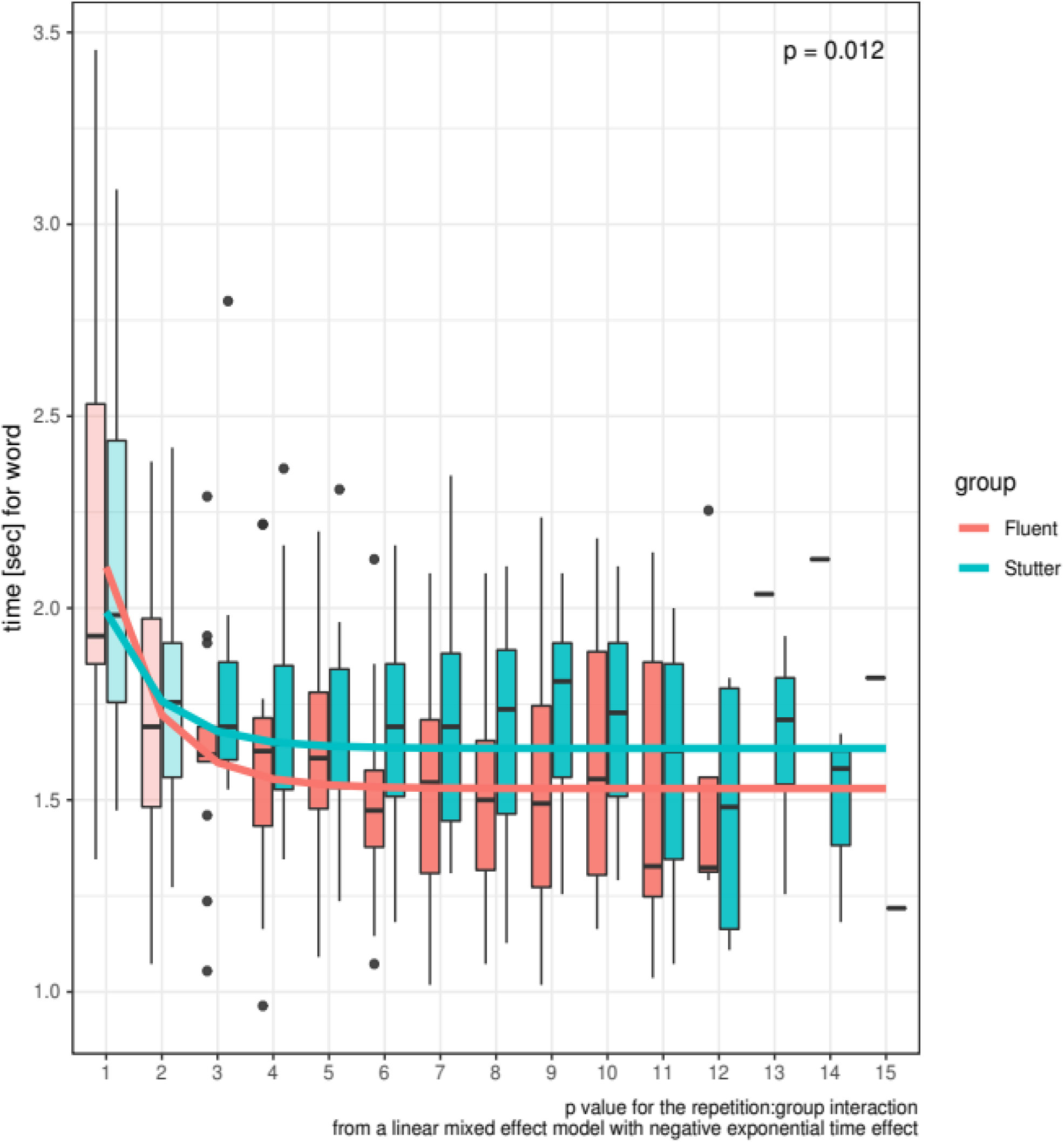
Speech duration. Time needed for speaking a pseudoword across the first ten correctly articulated and not stuttered repetitions. Displayed are box plots for each repetition in a group of adults who stutter and a group of control subjects. The lines show the model fits in both groups, the p value is from the interaction between repetition and group. For details see text.

## DISCUSSION

We used sophisticated analyses of real-time MRI image series capturing articulatory movements during fluent-sounding speech in adults who do and who do not stutter. We observed reproducible movement patterns of the inner and outer articulators which were similar in both groups.

Speech duration of the fluently spoken pseudoword “natscheitideut” was similar in both groups and decreased over repetitions, more so in ANS than in AWS.

The variability of the movement patterns of tongue, lips and velum decreased over repetitions. The extent of variability decrease was similar in both groups. Across all participants, this repetition effect on movement variability for the lips and the tip of the tongue was less pronounced in severely as compared to mildly stuttering individuals.

Typical stuttering is an intermittent phenomenon. A classical example is a pupil speaking fluently to his peers on the schoolyard, but stuttering severely in the classroom. This paroxysmal nature of stuttering generates misperceptions and increases the social burden of the disease (Blood et al. 2011).

Attempts to chart the border between fluency and dysfluent speech yielded opposing results. On the one hand, there is a discussion which types of dysfluencies can or cannot be termed stuttering. In children, some postulate that stuttering clearly differs from developmental dysfluencies (Sandrieser and Schneider 2008). In adults, the question what to count as stuttering events is vividly debated, in particular with regard to whole word repetitions (see online discussion of Reilly et al. 2009). This is supported by early studies from Johnson and Associates summarized in chapter 12 of the handbook (Bloodstein and Ratner 2008), which regard phrase repetitions, pauses and interjections as normal dysfluencies, and the syllable repetitions and sound prolongations as stuttering behaviors (page 307, Bloodstein and Ratner 2008).

At the other end of the spectrum, the continuity hypothesis postulates a gradual transition between fluency and dysfluency along a continuum (Adams and Runyan 1981).

The question per se has been challenged, because it is a mere question of definition whether subtle differences in adults who stutter are traces of stuttering, anticipation of future stuttering events or something different (see discussion on page 165 of Bloodstein and Ratner 2008). In addition, it is always difficult to prove that something does not exist. One can question the sensitivity of the methods employed to detect subtle differences in fluency that may exist beyond the listener’s ear.

In more general terms, the question arises to what extent motor deficits in stuttering generalize beyond the actual moments of dysfluency. If there is a disconnection of speech relevant brain areas (Sommer et al. 2002; Neef, Anwander, and Friederici 2015), it is always there and might translate into disabilities. According to the Packman and Attanasio model of stuttering, this chronic neurological disturbance per se is not sufficient to cause dysfluency, but only when accompanied by triggers such as linguistic complexity or variable syllabic stress. This interplay is assumed to be modulated by the level of arousal (Packman 2012), which impairs motor control (Yoshie et al. 2009). Essentially, their model postulates a threshold of the neurological imperfection to become detectable as stuttering. This is commonplace in neurology where disorders such as epilepsies are always present (and may be detectable by sophisticated electrophysiological means such as electroencephalography), but only intermittently clinically observable.

With its high time resolution, electroencephalography is indeed a useful tool to study state markers of dysfluency in stuttering, i.e. to understand what distinguishes stuttered from fluent utterances in PWS. Vanhoutte and colleagues (Vanhoutte et al. 2016) observed an enlarged contingent negative variation over right motor areas before fluent relative to stuttered utterances. They interpreted this as enhanced compensatory motor preparation over the right motor areas during fluent phases of speech.

This is consistent with an updated meta-analysis of fMRI studies, which yielded state-related excess of BOLD signal in the right (Belyk, Kraft, and Brown 2017) or bilateral primary motor cortex (Connally et al. 2018). Our findings support focussing research endeavours on state markers of dysfluency.

## Legends to tables and figures

**Supplementary table 1:** Comparison of variability index across articulation loci.

**Supplementary tables 2-4:** Variability of articulatory movements for different articulators, in early and late trials, and in adults who stutter and a group of control participants. For details see text.

**Supplementary figure 1:** Example spectogram: frequency in kHz (y axis) by time in seconds (x-axis) colored by energy level. Marks show the starting point (plosive [g]) and end point (plosive [b]). Dark area shows the segment for analysis.

**Supplementary figure 2:** The start and end points are checked visually and auditorily. The left panel shows an example of the start point at the plosive [g] at which the back of the tongue touches the velum. The right image shows the end point at the plosive [b] where the lips close.

**Supplementary figure 3:** Example of the initial grid definition. The left panel shows one single frame of the real-time MRI video. As reference line the upper front edge of the cervical vertebrae 4 and the transition from the anterior incisor to the hard palate were manually marked and connected (yellow line). The midpoint of the baseline was the starting point for seven lines drawn automatically at angles of 0 (yellow), 30 (pink), 60 (cyan), 90 (red), 120 (green), 150 (white), and 180 (yellow) degrees between the two halves of the baseline (supplementary figure 3). The right panel shows the extracted line profiles, where the pixel values of the individual lines of the grid were analyzed over the time of the real-time MRI scan.

**Supplementary figure 4:** In-house software to refine gap detection. The central image displays (from bottom to top) the extraction of the MRI video, a pixel map of the detected thin air, an overlay of the pixel map over the extracted MRI image, the measured gap width (in pixels). The left side bar shows the data selector, the right side bar shows the parameters that can be adjusted.

**Supplementary figure 5:** Examples of semi-automated gap detection and gap measurement in each of the 7 extracted dimension. In each image, the bottom panel shows the extracted line profile, the third panel show a pixel map of the detected thin air, the second panel shows an overlay of the pixel map over the extracted line profile, and the top panel shows the measured gap width (in pixels).

**Supplementary figures 6-12:** Modeled movement of articulators over time. The movement patterns were modeled using mixed-effect penalized functional regression models. Group-level model predictions for dimension 1-7s; thin lines = measured individual profiles, thick lines = group-level model predictions.

## 1 Ethics Statement

This study was carried out in accordance with the recommendations of the Ethics committee of the University Medical Center Göttingen with written informed consent from all subjects. All subjects gave written informed consent in accordance with the Declaration of Helsinki. The protocol was approved by the Ethics committee of the University Medical Center Göttingen.

## 2 Conflict of Interest

The authors declare that the research was conducted in the absence of any commercial or financial relationships that could be construed as a potential conflict of interest.

## 3 Author Contributions

MS designed the experiments, developed the analysis pipeline and wrote the first draft. AL developed the analysis pipeline and calculated the statistics. SD performed the file selection and film clip processing. DP recorded the data and initiated data analysis. AP contributed to study design and discussed the results. AK contributed to study design and analysis. WP provided funding and commented on the analysis and interpretation of the data. AJ performed the MRI experiments. JF developed the setup, contributed to the statistical analysis of the data, and helped writing the manuscript. All authors discussed the results and commented on the manuscript.

## 4 Funding

This work was supported by a seed grant from the primate cognition Leibniz-WissenschaftsCampus (Sommer/Frahm DM 22-606) to MS and JF

## 5 Acknowledgments

We are grateful to Bettina Helten for analyzing the speech samples and to Fabian Scheipl for detailed answers to questions on the functional regression.

## 6 Data Availability Statement

Pseudonymized data can be accessed by future researchers upon reasonable request based on standard hospital practices.

