## Supplementary Material for "Iceberg or cut off – how adults who stutter articulate fluent-sounding utterances"

Supplementary table 1

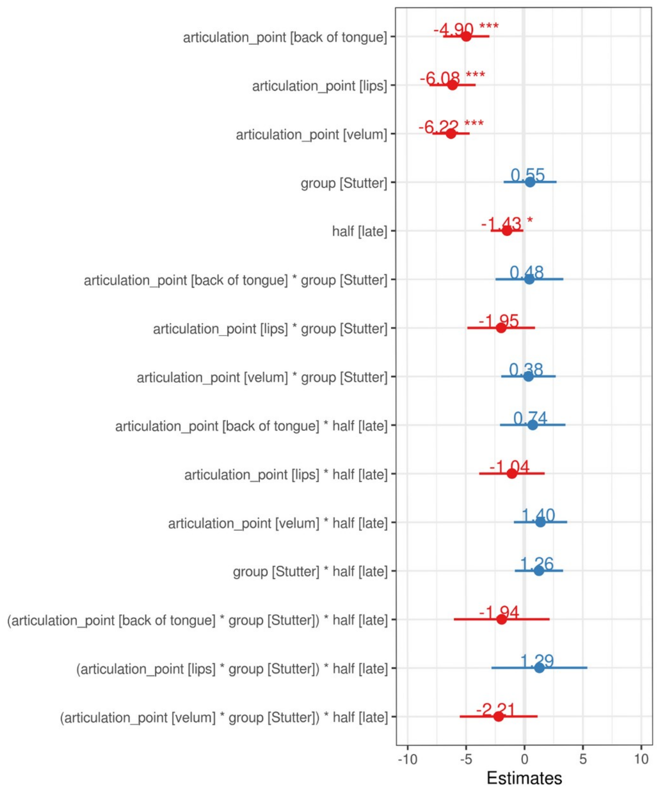


| term | estimate | ci | p.value |
|---|---|---|---|
| articulation_point [back of tongue] | -4.90 | [-6.88; -2.92] | < 0.001 *** |
| articulation_point [lips] | -6.08 | [-8.06; -4.10] | < 0.001 *** |
| articulation_point [velum] | -6.22 | [-7.81; -4.62] | < 0.001 *** |
| group [Stutter] | 0.55 | [-1.71; 2.81] | 0.633 |
| half [late] | -1.43 | [-2.83; -0.03] | 0.045 * |
| articulation_point [back of tongue] * group [Stutter] | 0.48 | [-2.42; 3.38] | 0.748 |
| articulation_point [lips] * group [Stutter] | -1.95 | [-4.85; 0.95] | 0.189 |
| articulation_point [velum] * group [Stutter] | 0.38 | [-1.95; 2.72] | 0.747 |
| articulation_point [back of tongue] * half [late] | 0.74 | [-2.06; 3.54] | 0.603 |
| articulation_point [lips] * half [late] | -1.04 | [-3.84; 1.76] | 0.465 |
| articulation_point [velum] * half [late] | 1.40 | [-0.88; 3.68] | 0.229 |
| group [Stutter] * half [late] | 1.26 | [-0.81; 3.33] | 0.233 |
| (articulation_point [back of tongue] * group [Stutter]) * half [late] | -1.94 | [-6.04; 2.16] | 0.355 |
| (articulation_point [lips] * group [Stutter]) * half [late] | 1.29 | [-2.81; 5.39] | 0.539 |
| (articulation_point [velum] * group [Stutter]) * half [late] | -2.21 | [-5.54; 1.12] | 0.194 |

Supplementary table 2

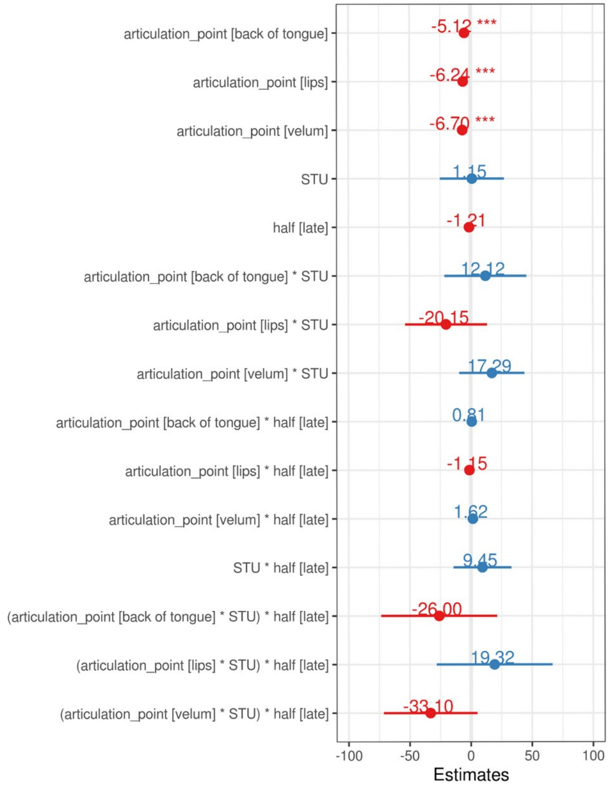


| term | estimate | ci | p.value |
|---|---|---|---|
| articulation_point [back of tongue] | -5.12 | [ -7.03; -3.22] | < 0.001 *** |
| articulation_point [lips] | -6.24 | [ -8.15; -4.34] | < 0.001 *** |
| articulation_point [velum] | -6.70 | [ -8.24; -5.15] | < 0.001 *** |
| STU | 1.15 | [-25.10; 27.41] | 0.931 |
| half [late] | -1.21 | [ -2.56; 0.15] | 0.080 |
| articulation_point [back of tongue] * STU | 12.12 | [-21.44; 45.67] | 0.479 |
| articulation_point [lips] * STU | -20.15 | [-53.71; 13.40] | 0.239 |
| articulation_point [velum] * STU | 17.29 | [ -9.41; 44.00] | 0.204 |
| articulation_point [back of tongue] * half [late] | 0.81 | [ -1.88; 3.51] | 0.554 |
| articulation_point [lips] * half [late] | -1.15 | [ -3.85; 1.55] | 0.405 |
| articulation_point [velum] * half [late] | 1.62 | [ -0.57; 3.82] | 0.147 |
| STU * half [late] | 9.45 | [-14.29; 33.19] | 0.435 |
| (articulation_point [back of tongue] * STU) * half [late] | -26.00 | [-73.45; 21.45] | 0.283 |
| (articulation_point [lips] * STU) * half [late] | 19.32 | [-28.13; 66.77] | 0.425 |
| (articulation_point [velum] * STU) * half [late] | -33.10 | [-71.39; 5.18] | 0.090 |

A

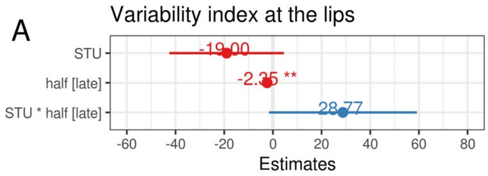


| term | estimate | ci | p.value | |
|---|---|---|---|---|
| STU | -19.0 | [-42.5; 4.54] | 0.114 | |
| half [late] | -2.4 | [ -4.1; -0.63] | 0.008 | ** |
| STU * half [late] | 28.8 | [ -1.6; 59.17] | 0.064 | |

B

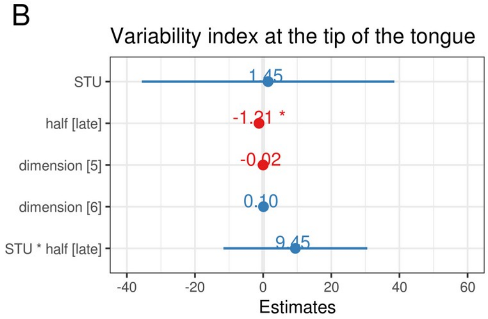


| term | estimate | ci | p.value | |
|---|---|---|---|---|
| STU | 1.45 | [-35.6; 38.5] | 0.939 | |
| half [late] | -1.21 | [ -2.4; 0.0] | 0.049 | * |
| dimension [5] | -0.02 | [ -1.1; 1.1] | 0.975 | |
| dimension [6] | 0.10 | [ -1.0; 1.2] | 0.868 | |
| STU * half [late] | 9.45 | [-11.7; 30.6] | 0.380 | |

C

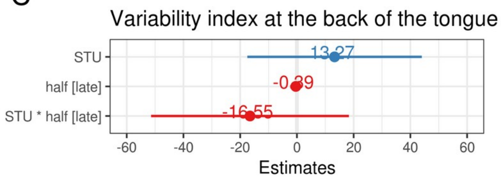


| term | estimate | ci | p.value |
|---|---|---|---|
| STU | 13.27 | [-17.5; 44.0] | 0.398 |
| half [late] | -0.39 | [ -2.4; 1.6] | 0.699 |
| STU * half [late] | -16.55 | [-51.4; 18.3] | 0.352 |

D

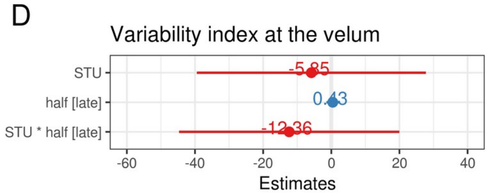


| term | estimate | ci | p.value |
|---|---|---|---|
| STU | -5.85 | [-39.5; 27.8] | 0.733 |
| half [late] | 0.43 | [ -1.4; 2.3] | 0.653 |
| STU * half [late] | -12.36 | [-44.7; 20.0] | 0.454 |

Supplementary table 3

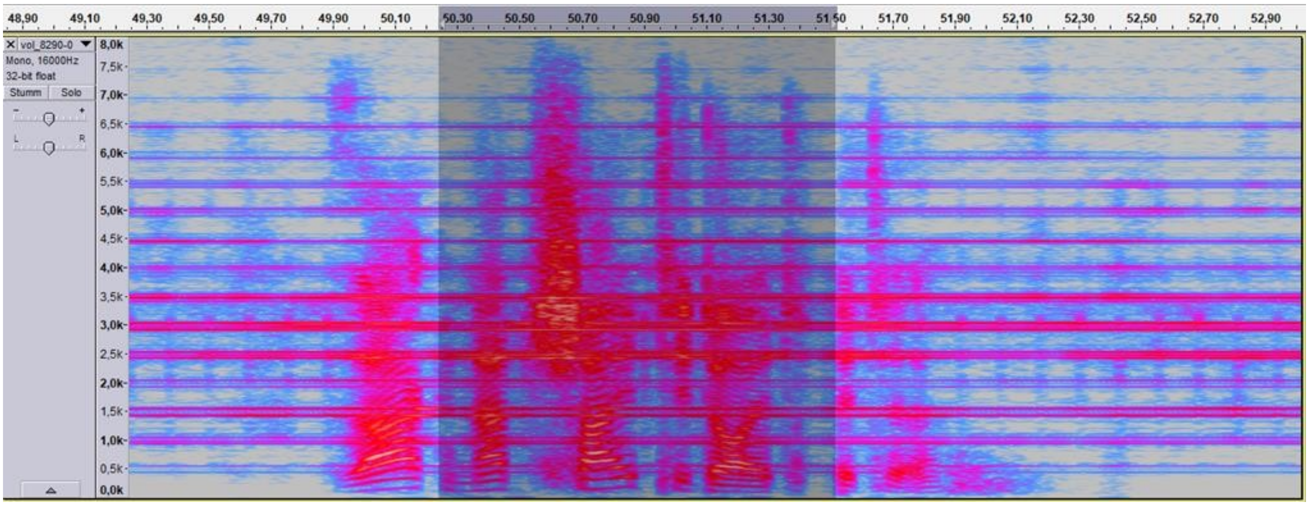


Supplementary figure 1

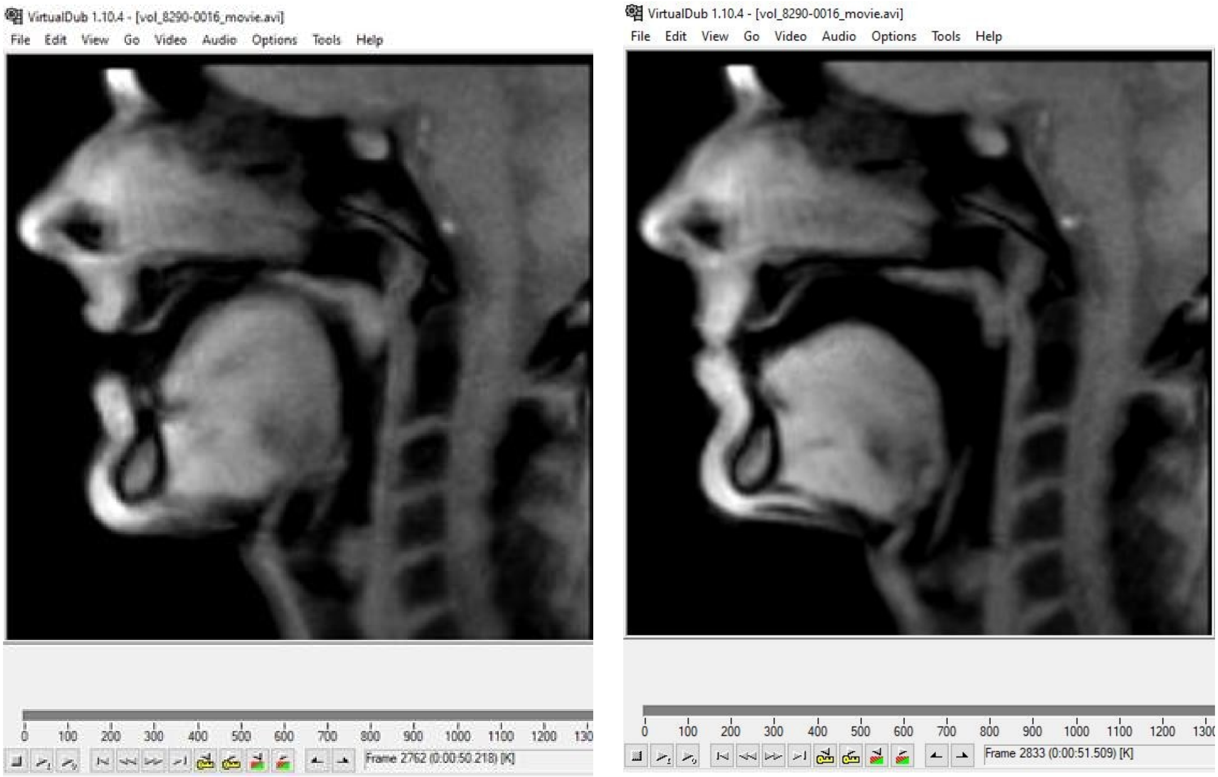


Supplementary figure 2

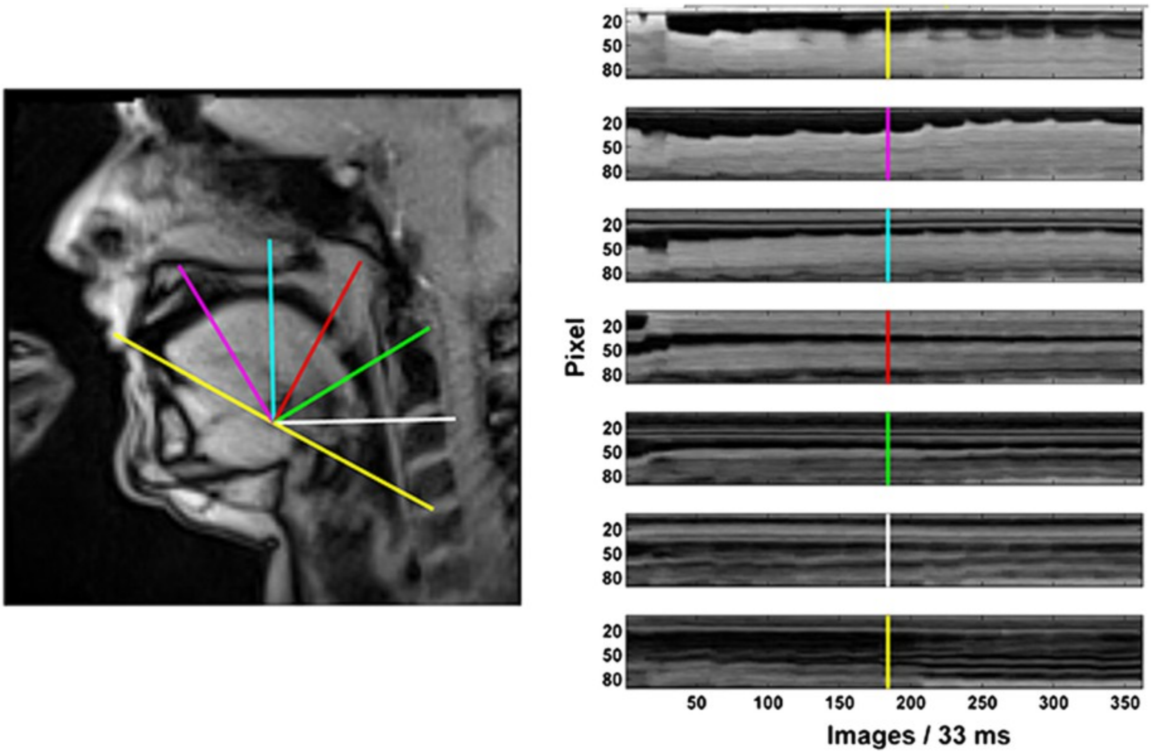


Supplementary figure 3

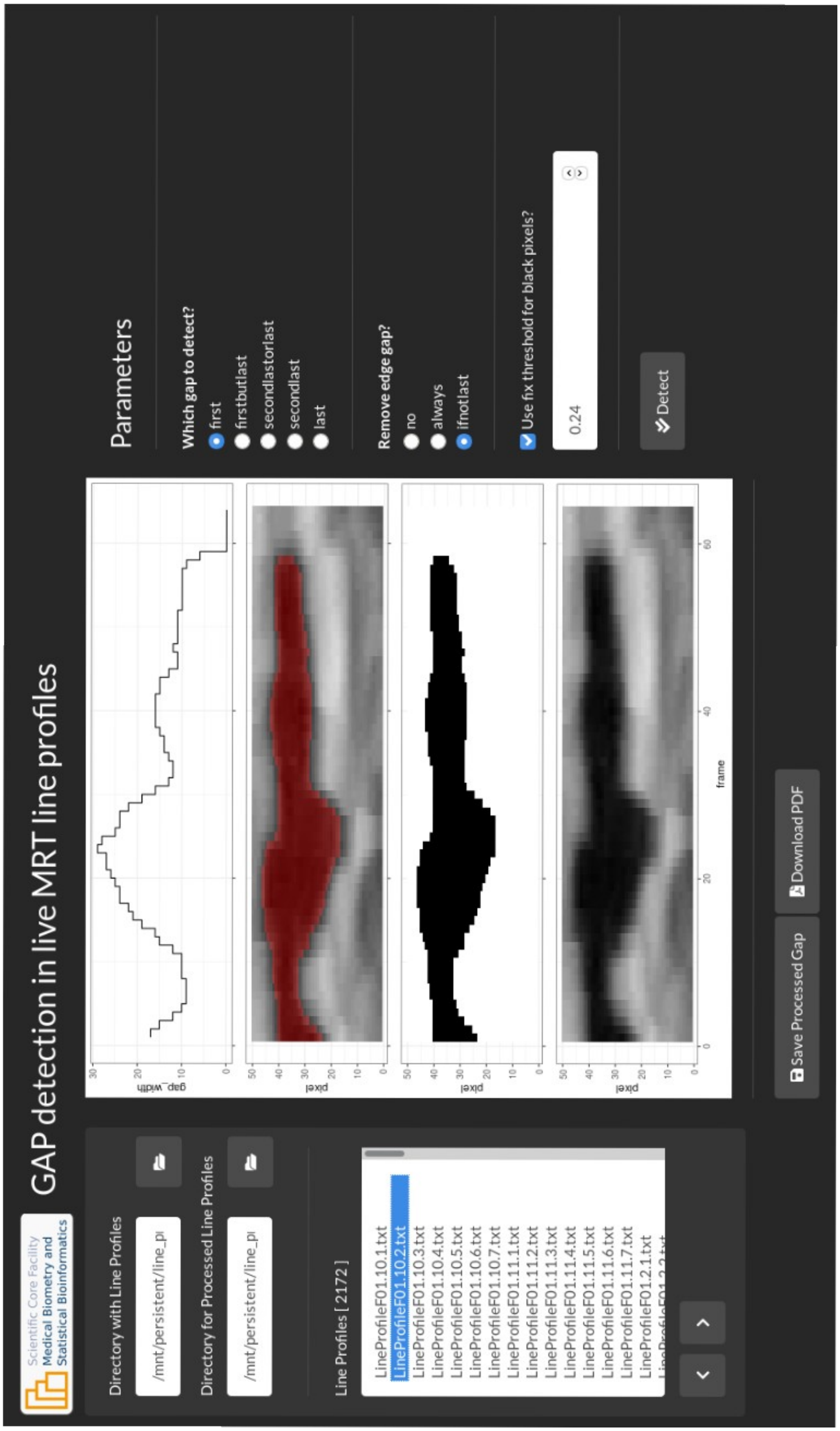


Supplementary figure 4

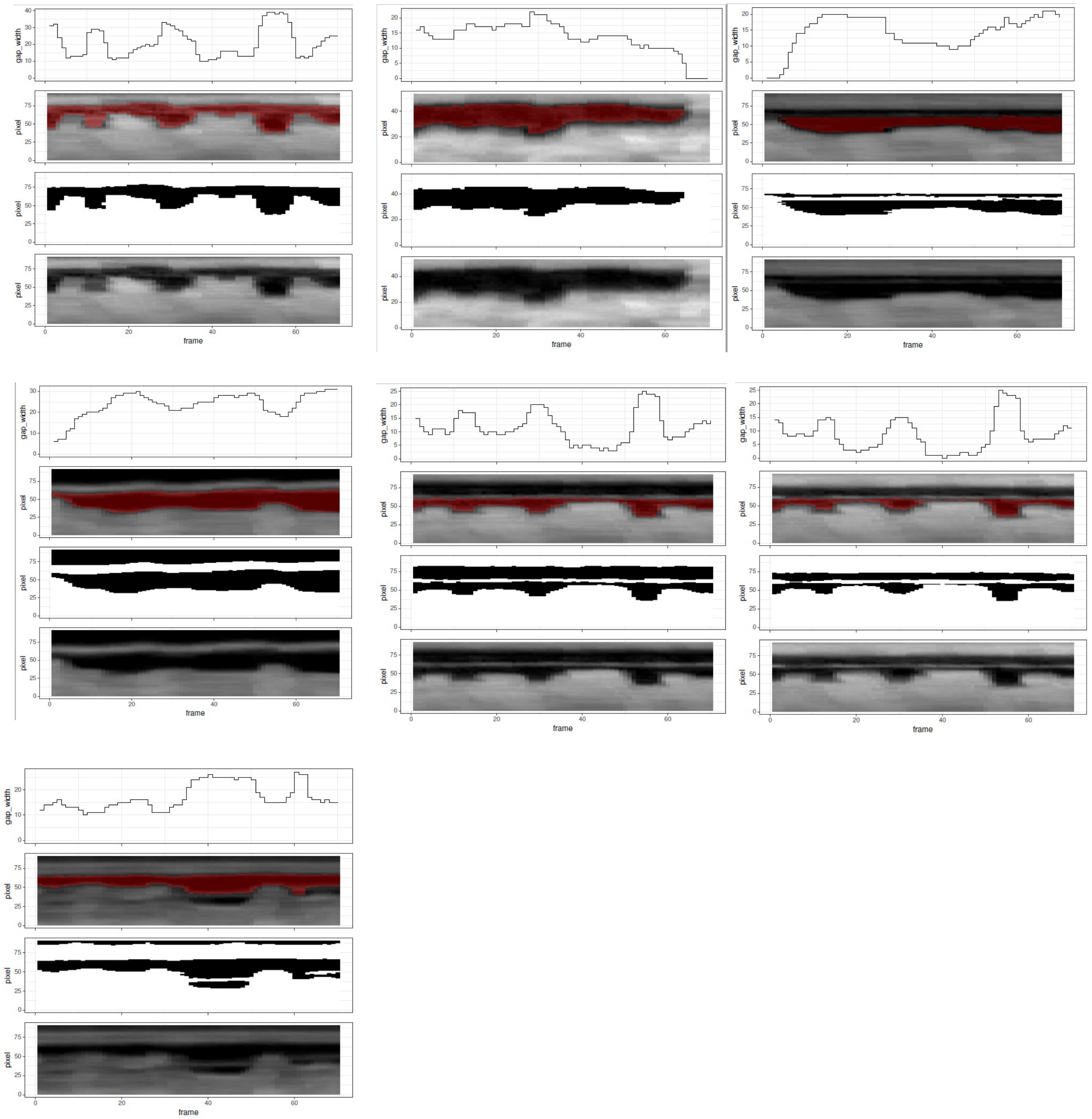


Supplementary figure 5

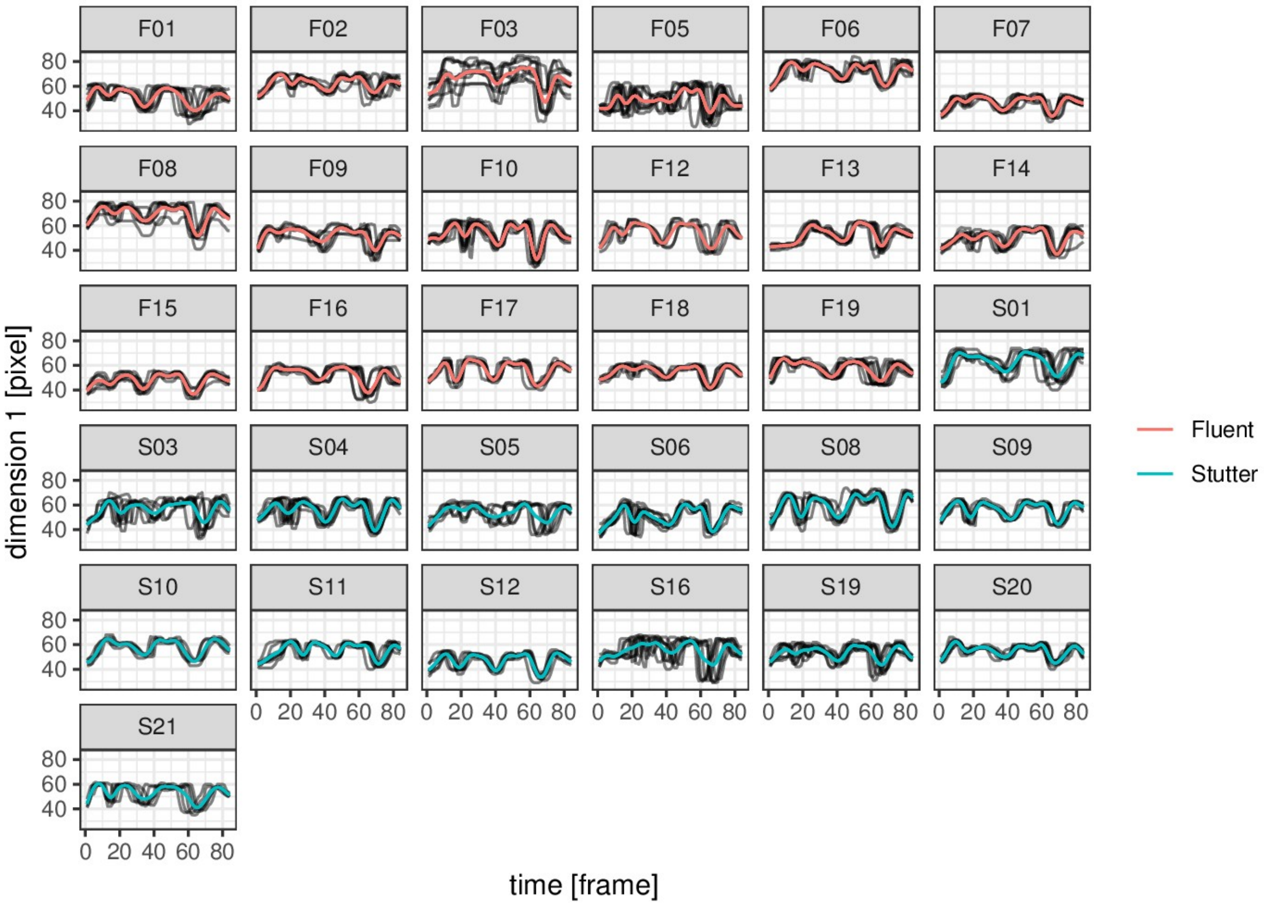


Supplementary figure 6

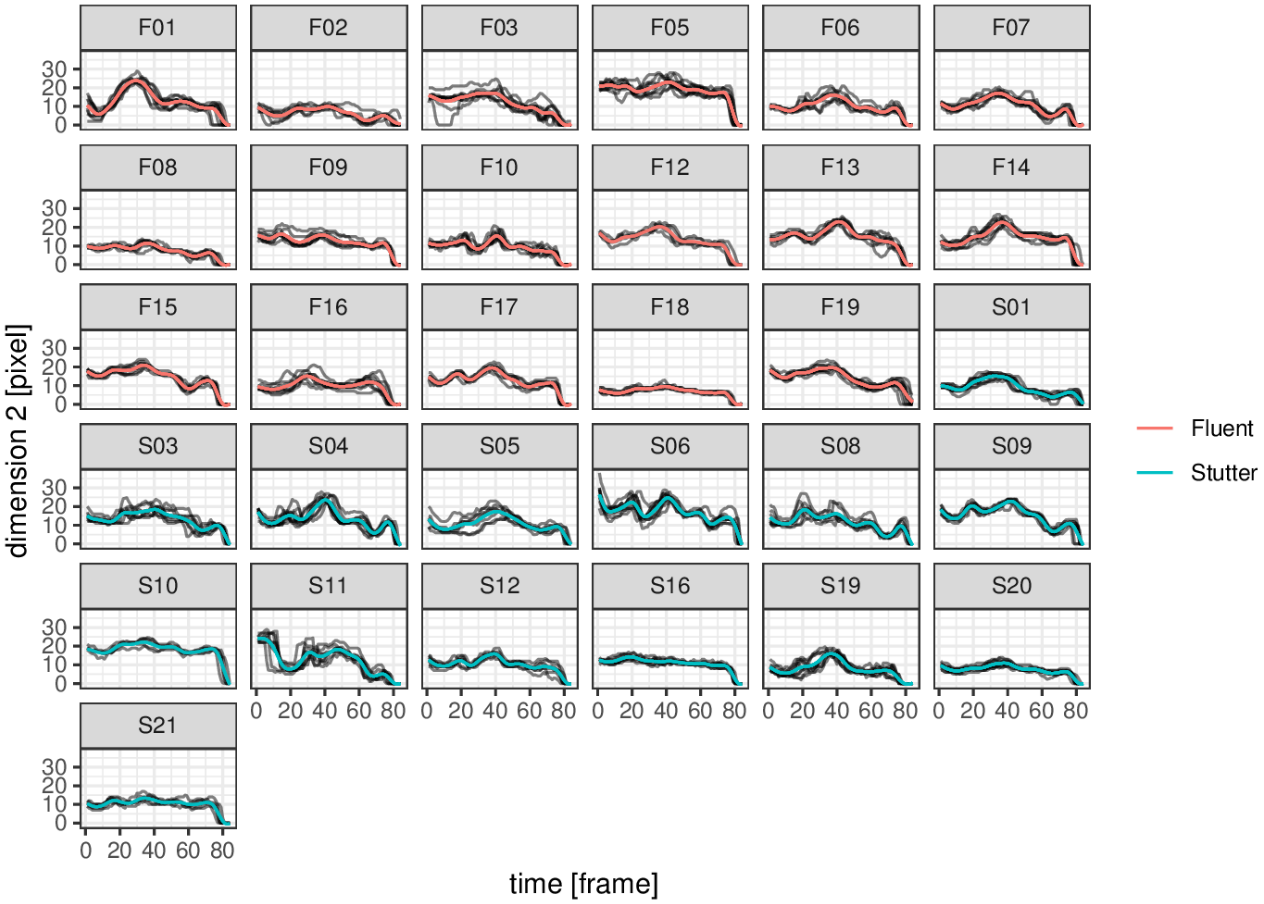


Supplementary figure 7

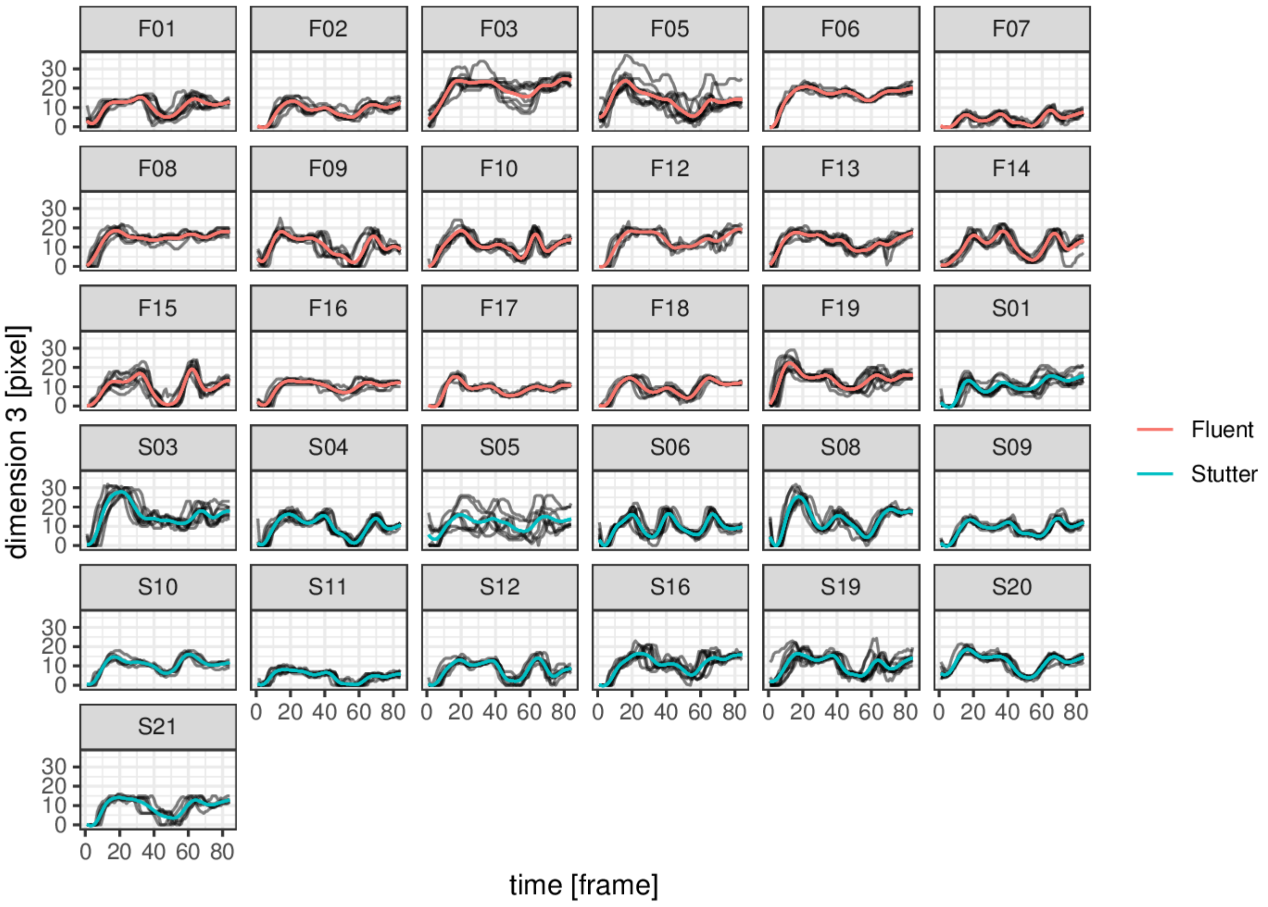


Supplementary figure 8

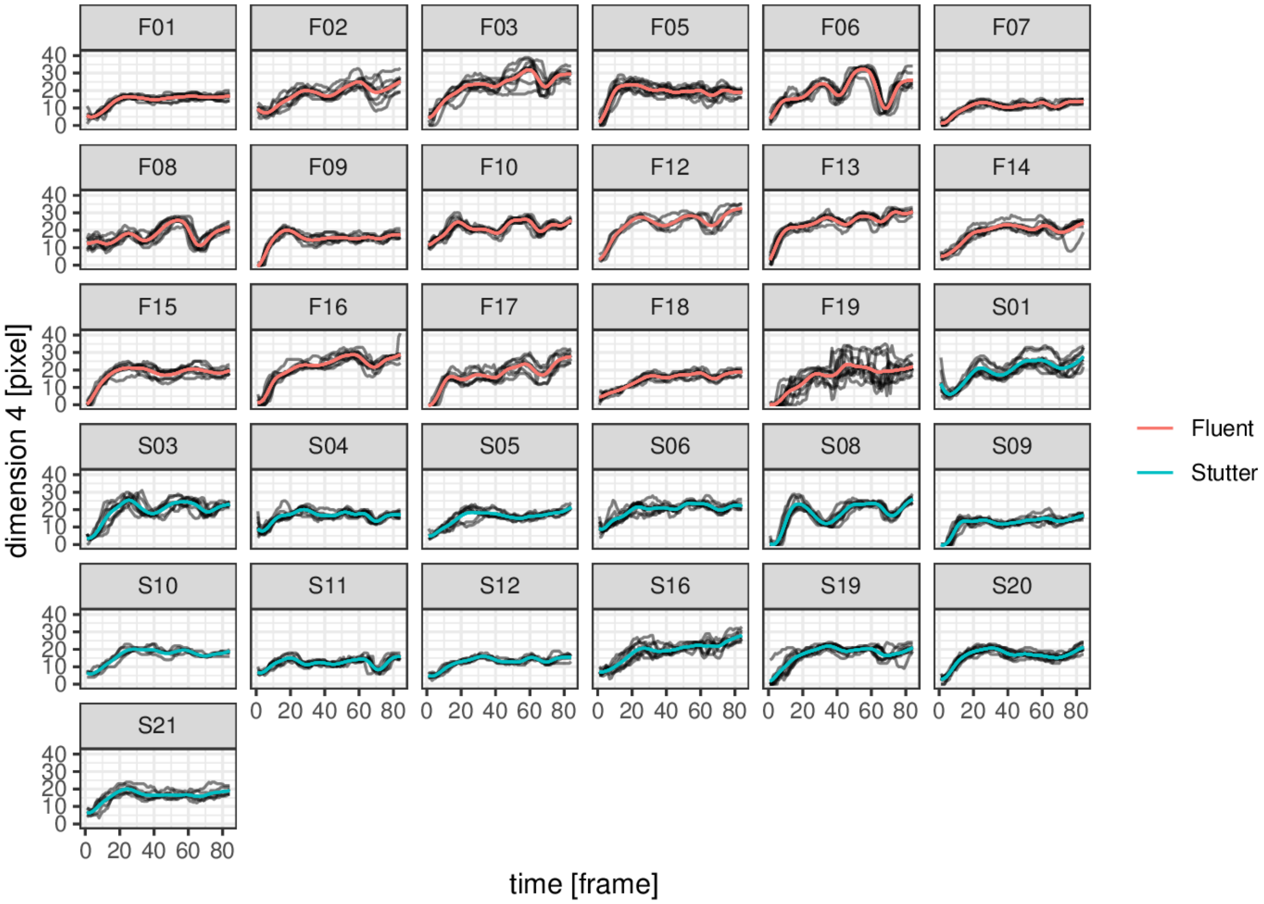


Supplementary figure 9

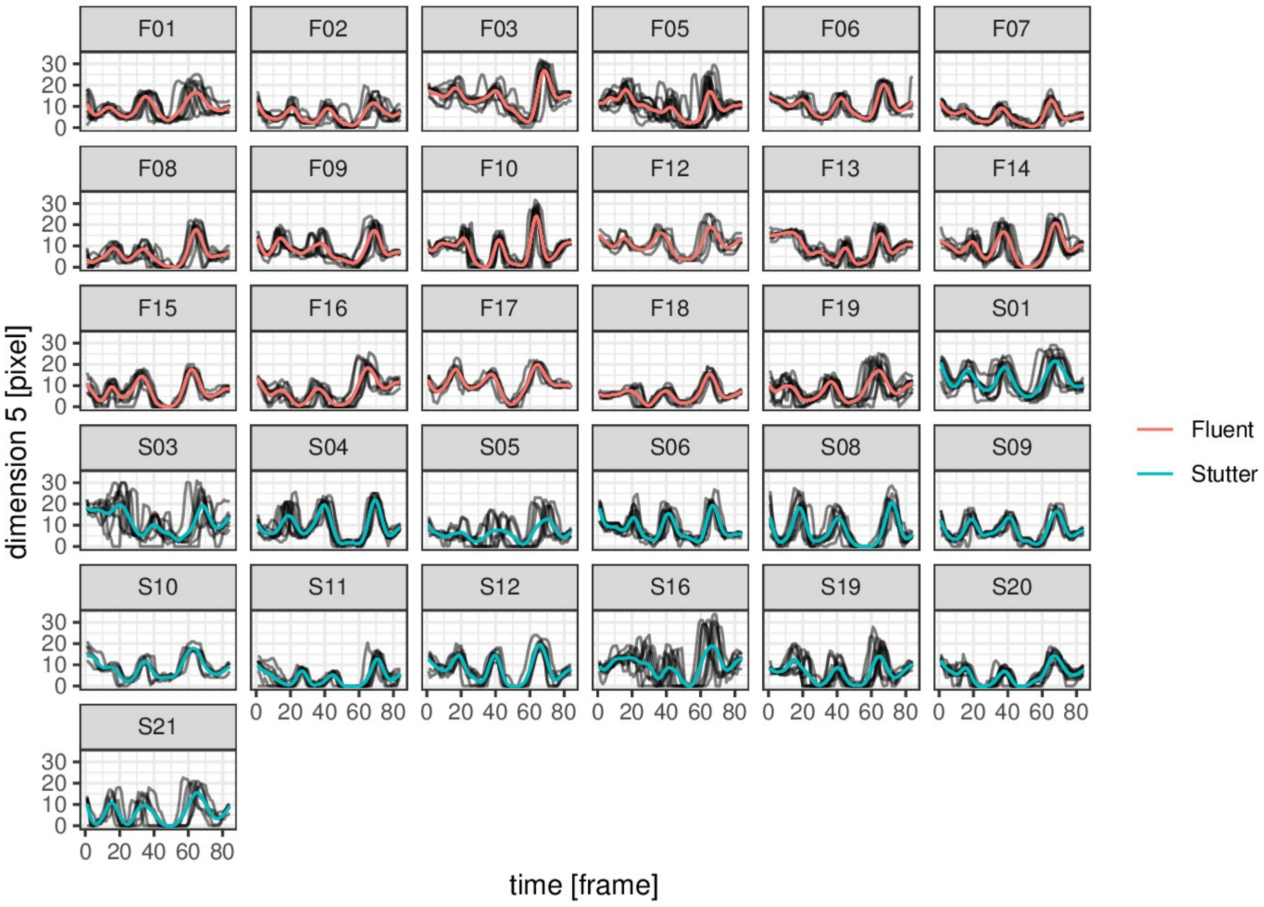


Supplementary figure 10

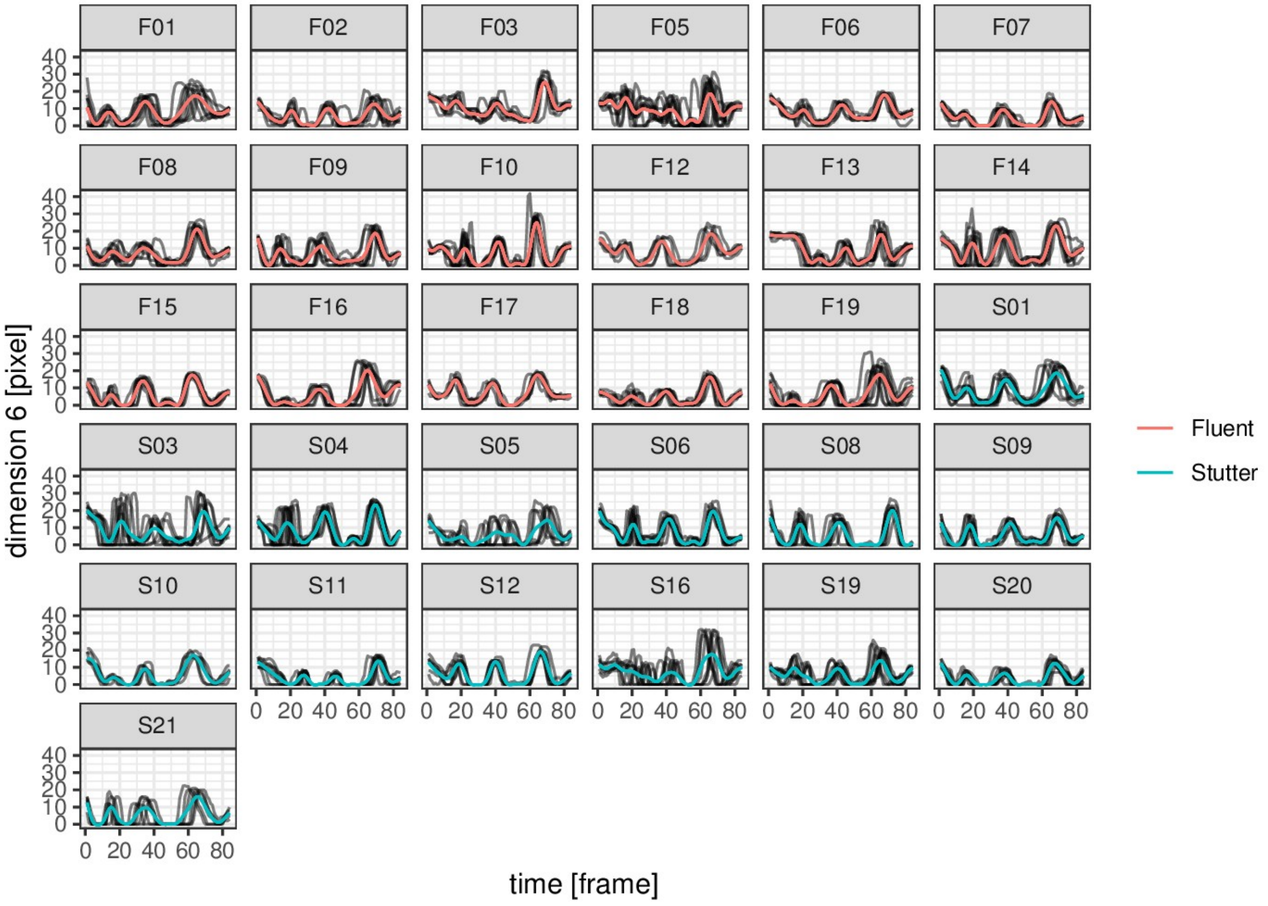


Supplementary figure 11

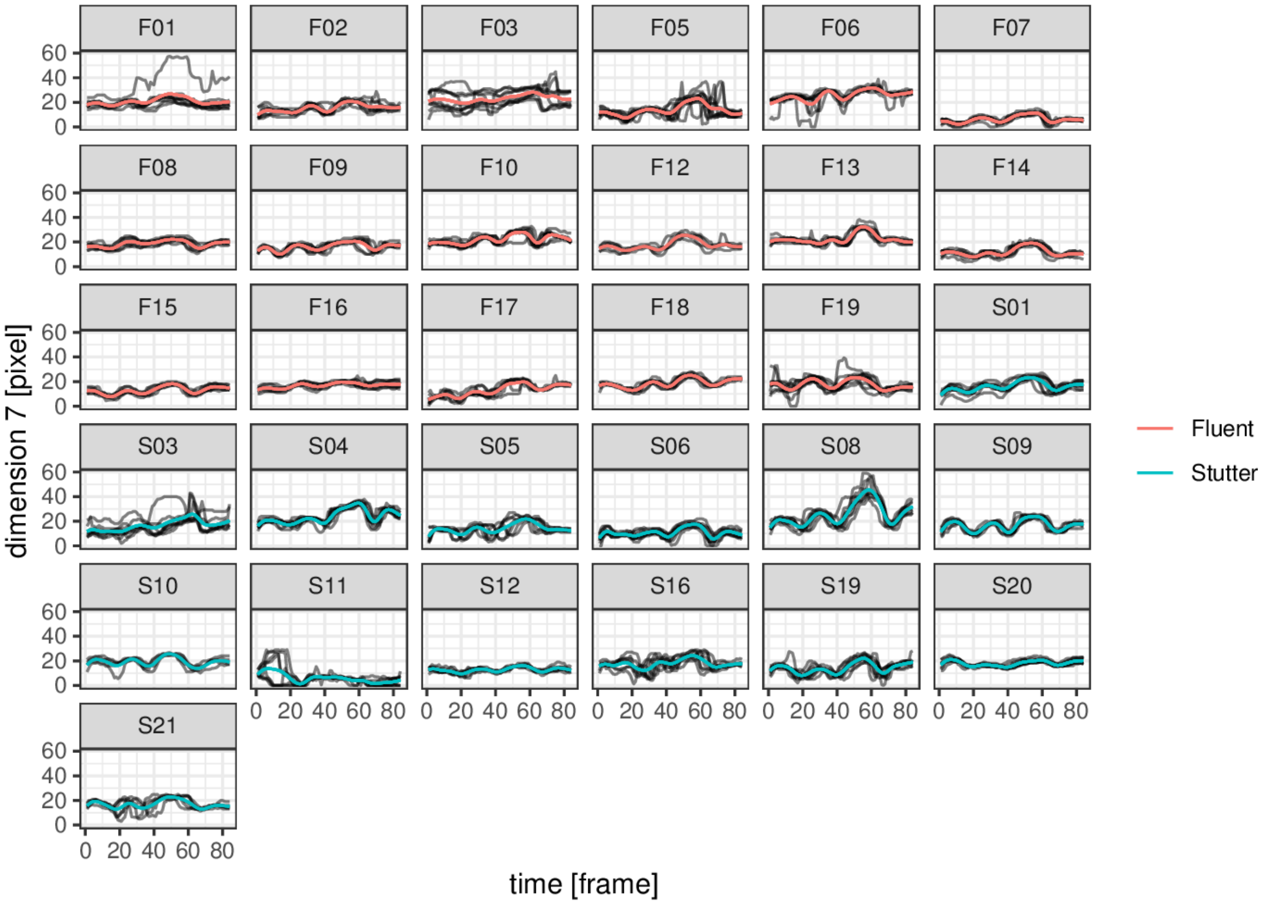


Supplementary figure 12

Dimension 1

Constant coefficients:

| term | Estimate | Std. Error | t value | Pr(>\|t\|) |
|---|---|---|---|---|
| (Intercept) | 56.0 | 0.45 | 123.0 | < 0.001 |

Smooth terms & functional coefficients:

| term | edf | Ref.df | F | p-value |
|---|---|---|---|---|
| Intercept(yindex) | 17.0 | 19.0 | 12.01 | < 0.001 |
| group(yindex) | 1.0 | 1.0 | 0.71 | 0.399 |
| s(proband) | 628.0 | 773.0 | 55.17 | < 0.001 |

Dimension 2

Constant coefficients:

| term | Estimate | Std. Error | t value | Pr(>\|t\|) |
|---|---|---|---|---|
| (Intercept) | 12.0 | 0.29 | 40.0 | < 0.001 |

Smooth terms & functional coefficients:

| term | edf | Ref.df | F | p-value |
|---|---|---|---|---|
| Intercept(yindex) | 17.0 | 19.0 | 15.1 | < 0.001 |
| group(yindex) | 19.0 | 20.0 | 3.8 | < 0.001 |
| s(proband) | 534.0 | 773.0 | 80.8 | < 0.001 |

Dimension 3

Constant coefficients:

| term | Estimate | Std. Error | t value | Pr(>\|t\|) |
|---|---|---|---|---|
| (Intercept) | 11.0 | 0.29 | 38.0 | < 0.001 |

Smooth terms & functional coefficients:

| term | edf | Ref.df | F | p-value |
|---|---|---|---|---|
| Intercept(yindex) | 17.3 | 19.0 | 21.65 | < 0.001 |
| group(yindex) | 1.8 | 1.9 | 0.43 | 0.585 |
| s(proband) | 551.8 | 773.0 | 58.84 | < 0.001 |

Dimension 4

Constant coefficients:

| term | Estimate | Std. Error | t value | Pr(>\|t\|) |
|---|---|---|---|---|
| (Intercept) | 17.0 | 0.32 | 54.0 | < 0.001 |

Smooth terms & functional coefficients:

| term | edf | Ref.df | F | p-value |
|---|---|---|---|---|
| Intercept(yindex) | 17.0 | 19.0 | 17.0 | < 0.001 |
| group(yindex) | 1.0 | 1.0 | 6.0 | 0.014 |
| s(proband) | 506.0 | 773.0 | 63.0 | < 0.001 |

Dimension 5

Constant coefficients:

| term | Estimate | Std. Error | t value | Pr(>\|t\|) |
|---|---|---|---|---|
| (Intercept) | 8.30 | 0.23 | 37.0 | < 0.001 |

Smooth terms & functional coefficients:

| term | edf | Ref.df | F | p-value |
|---|---|---|---|---|
| Intercept(yindex) | 17.2 | 19.0 | 18.3 | < 0.001 |
| group(yindex) | 1.1 | 1.1 | 1.9 | 0.180 |
| s(proband) | 622.5 | 773.0 | 29.6 | < 0.001 |

Dimension 6

Constant coefficients:

| term | Estimate | Std. Error | t value | Pr(>\|t\|) |
|---|---|---|---|---|
| (Intercept) | 6.50 | 0.21 | 31.0 | < 0.001 |

Smooth terms & functional coefficients:

| term | edf | Ref.df | F | p-value |
|---|---|---|---|---|
| Intercept(yindex) | 17.0 | 19.0 | 23.6 | < 0.001 |
| group(yindex) | 1.0 | 1.0 | 2.8 | 0.095 |
| s(proband) | 644.0 | 773.0 | 22.6 | < 0.001 |

Dimension 7

Constant coefficients:

| term | Estimate | Std. Error | t value | Pr(>\|t\|) |
|---|---|---|---|---|
| (Intercept) | 17.0 | 0.44 | 39.0 | < 0.001 |

Smooth terms & functional coefficients:

Supplementary table 4
